# Interplay of Monocytes and T Lymphocytes in COVID-19 Severity

**DOI:** 10.1101/2020.07.17.209304

**Authors:** Lindsey E. Padgett, Huy Q. Dinh, Serena J. Chee, Claire E. Olingy, Runpei Wu, Daniel J. Araujo, Pandurangan Vijayanand, Christian H. Ottensmeier, Catherine C. Hedrick

## Abstract

The COVID-19 pandemic represents an ongoing global crisis that has already impacted over 13 million people. The responses of specific immune cell populations to the disease remain poorly defined, which hinders improvements in treatment and care management. Here, we utilized mass cytometry (CyTOF) to thoroughly phenotype peripheral myeloid cells and T lymphocytes from 30 convalescent patients with mild, moderate, and severe cases of COVID-19. We identified 10 clusters of monocytes and dendritic cells and 17 clusters of T cells. Examination of these clusters revealed that both CD14^+^CD16^+^ intermediate and CD14^dim^CD16^+^ nonclassical monocytes, as well as CD4^+^ stem cell memory T (T_SCM_) cells, correlated with COVID-19 severity, coagulation factor levels, and/or inflammatory indicators. We also identified two nonclassical monocyte subsets distinguished by expression of the sugar residue 6-Sulfo LacNac (Slan). One of these subsets (Slan^lo^, nMo1) was depleted in moderately and severely ill patients, while the other (Slan^hi^, nMo2) increased with disease severity and was linked to CD4^+^ T effector memory (T_EM_) cell frequencies, coagulation factors, and inflammatory indicators. Intermediate monocytes tightly correlated with loss of naive T cells as well as an increased abundance of effector memory T cells expressing the exhaustion marker PD-1. Our data suggest that both intermediate and non-classical monocyte subsets shape the adaptive immune response to SARS-CoV-2. In summary, our study provides both broad and in-depth characterization of immune cell phenotypes in response to COVID-19 and suggests functional interactions between distinct cell types during the disease.

**One Sentence Summary:** Use of mass cytometry on peripheral blood mononuclear cells from convalescent COVID-19 patients allows correlation of distinct monocyte and T lymphocyte subsets with clinical factors.

## INTRODUCTION

The coronavirus disease 2019 (COVID-19) pandemic, caused by SARS-Coronavirus-2 (SARS-CoV-2), has currently affected more than 13 million people and claimed over 580,000 lives. Clinical presentation of COVID-19 varies, with some individuals exhibiting mild to severe symptoms while others remain asymptomatic. Severe cases are associated with acute respiratory distress syndrome, which requires hospitalization and mechanically-assisted ventilation (*1*). COVID-19 is also generally linked to an increased risk of cardiovascular events such as thrombosis (*1–4*), pulmonary embolism (*3, 4*), and stroke (*3, 4*), especially in critically ill patients. Comorbidities such as diabetes, hypertension, respiratory illness, and age (> 65 years) are common in severe cases (*1, 5–9*) and worsen prognosis (*5, 8, 10*). Improved treatment of COVID-19 therefore requires a refined understanding of the heterogeneity in immune responses to SARS-CoV-2 infection and the cellular subsets that drive these processes.

Peripheral inflammation is linked to development of severe COVID-19 (*11*), and secondary immune mechanisms may regulate progression of the disease (*12*). Monocytes are myeloid cells that govern innate immune responses to viral infection and have traditionally been classified as CD14^+^CD16^-^ (classical), CD14^lo^CD16^+^ (nonclassical), and CD14^+^CD16^+^ (intermediate). Expansion of CD14^+^HLA-DR^lo^, CD14^+^CD16^+^, and immunosuppressive, immature monocytes have been demonstrated in peripheral blood mononuclear cells (PBMCs) from severely ill COVID-19 patients (*11, 13, 14*). Patients with mild forms of COVID-19 display an increased abundance of CD14^+^HLA-DR^hi^CD11c^hi^ inflammatory monocytes (*14*). T cells are lymphocytes that coordinate adaptive immune responses to viral insults and bifurcate into CD4^+^ (helper) and CD8^+^ (cytotoxic) subsets. Activation of peripheral CD8^+^ and CD4^+^ T cell subsets is heterogeneous amongst COVID-19 patients (*15*). Circulating T cell numbers are also broadly reduced in moderate to severe cases, with CD8^+^ T cells being particularly affected (*15–17*). Still, as there are conflicting reports on the expression of exhaustion markers on T cells during the course of the disease (*15, 18–20*), the events controlling this depletion are unknown. In-depth profiling of myeloid and T cell populations from COVID-19 patients could allow for interrogation of these processes, identification of other cellular subsets altered during illness, and correlation of cellular phenotypes with clinical parameters.

Herein, we utilize cytometry by time-of-flight (CyTOF) to assess myeloid and T cell heterogeneity in PBMCs from mildly to severely ill COVID-19 patients. We used these data to characterize the interplay between these two cell types in disease severity. Specifically, we identify unique subpopulations of monocytes and T cells and find that their relative abundances are differentially correlated to disease intensity and clinical factors. In summary, our work highlights perturbations in both monocytes and T lymphocytes as a response to COVID-19 and suggests interactions between these two immune cell types during the disease.

## RESULTS

### Global Changes in Immune Cell Composition and Responses in COVID-19

PBMCs were obtained from the blood of 30 COVID-19 subjects during convalescence and analyzed using mass cytometry (Fig. 1A). Mildly ill patients (n = 8) were healthcare workers who tested positive for SARS-CoV-2 and became ill with COVID-19, but did not need hospitalization. The moderate (n = 13) and severe (n = 9) COVID-19 patient groups required hospitalization, with severely ill patients specifically requiring admittance to an intensive treatment unit (Fig. 1B). Patient demographics are detailed in Methods. Clinical data were obtained for moderately and severely ill patients while in hospital, and all patients in both groups presented with a clinically-defined inflammatory syndrome. We have limited clinical information on mildly ill COVID-19 individuals, as they were not hospitalized. Thus, the focus of our study was to determine what immune system perturbations distinguish moderate and severe cases of COVID-19.

**Figure 1.**
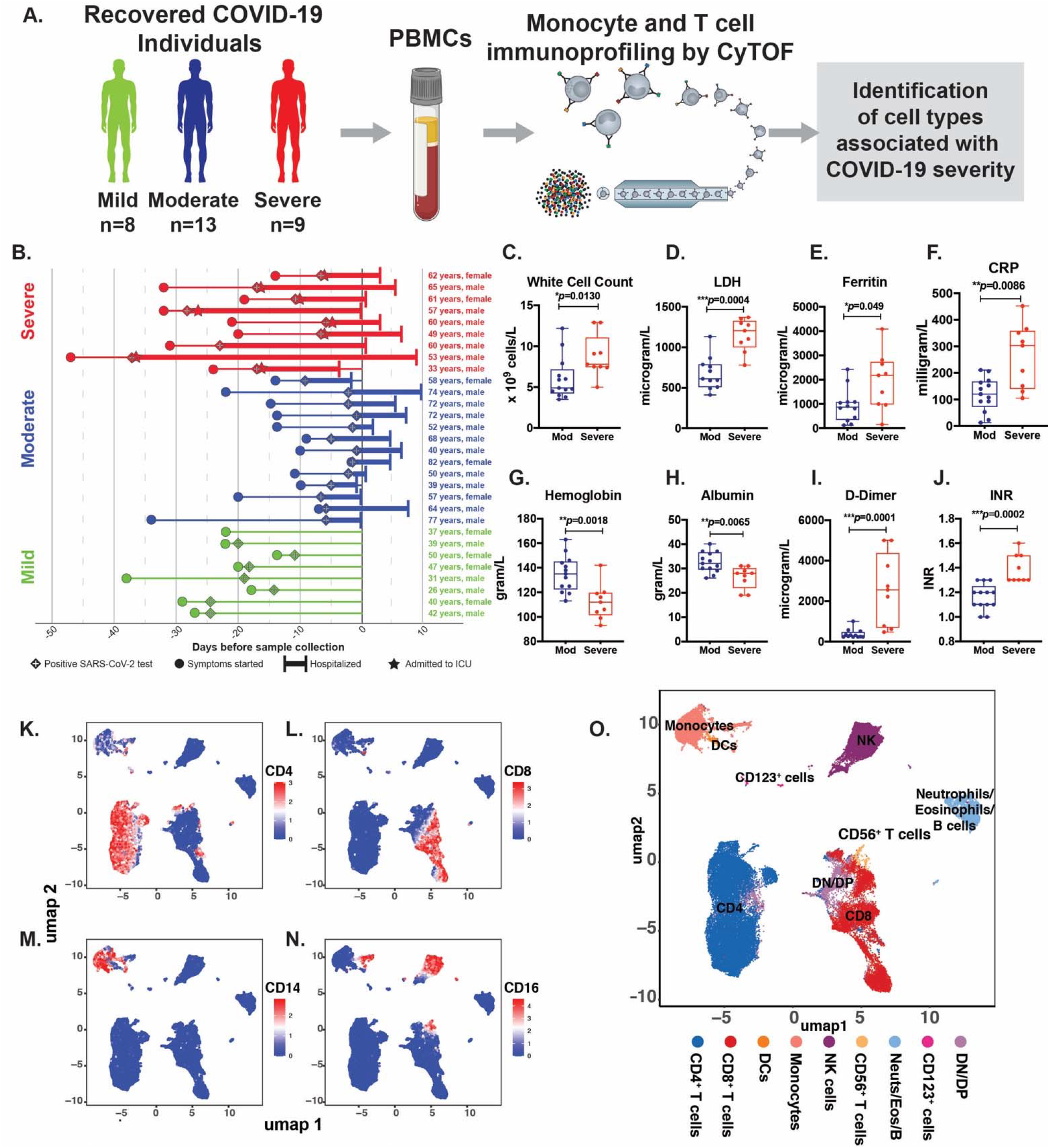
Experimental schematic for profiling monocyte and T cell phenotypes within recovered COVID-19 individuals. Monocyte and T cell immunoprofiling was performed via mass cytometry within PBMCs from mild (⊓=8), moderate (n=13), and severe (n=9) individuals (A). Patient cohort overview, depicting age, sex, COVID-19 severity, hospitalization, with all individuals aligned to the time of sample collection (**B**). Box and whisker plots of white blood cell counts (**C**), lactate dehydrogenase (LDH, **D**), Ferritin (**E**), C-Reactive Protein (CRP, **F**), hemoglobin (**G**), albumin (**H**), D-dimer (I), and international normalized ratio (INR, J) within moderate (n=13) and severe (n=8) individuals. CD45^+^ leukocytes were clustered and projected onto a UMAP, and expression of CD4 (**K**), CD8 (L), CD14 (**M**), and CD16 (**N**) were examined, leading to 9 immune populations (0). Statistical siginificance was calculated using Mann-whitney test.

Severely ill patients showed statistically significant increases in total white cell counts (Fig. 1C; *\*p*=0.013), lactate dehydrogenase (LDH, Fig. 1D; *\*\*\*p*=0.0004), ferritin (Fig. 1E; *\*p=0.049*), and C-Reactive Protein (CRP; *\*\*p=0*.0086; Fig. 1F) compared to moderately ill patients. Both hemoglobin (Fig. 1G; *\*\*p*=0.0018) and albumin (Fig. 1H; *\*\*p*=0.0065) levels were lower in severe disease. Severely affected patients also had higher plasma levels of D-dimers (Fig. 1I; *\*\*\*p*=0.0001) and international normalized ratio (INR) calculated from prothrombin time (Fig. 1J; *\*\*\*p*=0.0002). Platelet counts were also increased in severe disease (fig. S1A). While these latter data on platelet counts did not reach statistical significance, like D-dimers and INR, they are consistent with reports of increased coagulation events in COVID-19 patients (*9*). Oxygen saturation levels were also significantly reduced in severely ill patients (fig. S1B; *\*\*p*=0.003). Neutrophil:lymphocyte and neutrophil:monocyte ratios did not change, even though there was a trend towards increased neutrophil counts in severely ill patients (fig. S1C-D). Similarly, monocyte:lymphocyte ratios remained unchanged (fig. S1E). Comorbidities in hospitalized COVID-19 patients were primarily cardiovascular in scope, and included prior history of myocardial infarction and hypertension (see Methods).

Using a 41-marker CyTOF panel focused on identifying monocyte and T lymphocyte changes in COVID-19 (fig. S1F), we first assessed global immune cell changes in patients stratified by illness severity. We then utilized FlowSOM, an unbiased clustering method (*21*), along with meta-clustering and consensus clustering (see Methods) to identify 9 major immune cell types projected onto a Uniform Manifold Approximation and Projection plot (UMAP) plot (*22*), via assessment of markers for conventional T and monocyte cell types (Fig. 1K-O; fig. S2). UMAPs of the expression levels of all markers used to identify each CD45^+^ immune cell subset can be found in Supplementary Figure 2. Frequencies of all major immune cell types identified within the CD45^+^ gate for mildly (green), moderately (blue), and severely (red) ill COVID-19 patients are shown (fig. S1G,H). Specifically, we identified CD8^+^, CD4^+^, as well as double positive/negative (DN/DP), and CD56^+^ cells in the T cell compartment (fig. S1G; Fig. 1K,L; fig. S2). Other major cell types were dendritic cells (DC), natural killer (NK) cells, CD123^+^ cells (likely plasmacytoid dendritic cells (pDCs)) and monocytes (fig. S1H) using marker expression combinations of CD14 (Fig. 1M), CD16 (Fig. 1N), HLA-DR, CD1c, and CD123 (fig. S2). As our study was focused on monocyte/DC and T cells, markers for eosinophils, neutrophils, and B cells present in PBMC were placed in the same channel for exclusion (CD19/Siglec8/CD66b). We observed a slight reduction in CD4^+^ and CD8^+^ T cell frequencies in severe COVID patients, which is consistent with previous reports (*15, 16, 23–26*) (fig. S1G). We also found a significant increase in total monocyte frequencies stratified by disease severity, and a similar trend for CD1c^+^CD11c^hi^ DCs (fig. S1H).

### Classical Monocyte Heterogeneity in COVID-19

Heterogeneous myeloid cell phenotypes have been observed in COVID-19 patients (*13*), but how these phenotypes are dictated by SARS-CoV-2 illness is unknown. We designed a CyTOF panel to examine monocyte and DC heterogeneity in-depth, as we have done for our work in profiling immune cells during atherosclerosis (*27*). All monocytes and DCs identified in Figure 1C were re-clustered and projected onto a new UMAP to obtain a detailed myeloid heterogeneity landscape (Fig. 2A; fig. S3). Using FlowSOM clustering and meta-clustering to merge subsets of these myeloid cells, we identified 10 clusters containing monocytes and DCs in our COVID-19 cohort. These included 1 DC cluster, four CD14^+^ classical monocyte (cMo) clusters, 1 CD14^+^CD16^+^ intermediate monocyte (iMo) cluster, and 2 CD14^lo^CD16^+^ nonclassical monocyte (nMo) clusters (Fig. 2A). We also observed 2 new monocyte subsets present in blood of patients with COVID-19: CD14^+^CD16^+^CD56^+^ (CD56^+^) monocytes and CD14^+^CD3^+^ (CD3^+^) monocytes (Fig. 2A/C). CD56^+^ monocytes possess cytotoxic NK-like characteristics and have been identified in mice (*28*). A related CD14^+^CD56^+^ subset that lacks CD16 and is highly phagocytic has been reported in humans (*29*). CD3^+^ monocytes were also identified, and we confirmed that these were not doublets or cell aggregates (data not shown). Expression of CyTOF markers present on each myeloid cell cluster is shown in a heatmap in Figure 2B. We also observed that enrichment of each cluster varied between patients; however, all patients possessed all clusters (Fig. 2C). The classical monocyte cluster cMo1 increased in both moderate and severe cases compared to mild cases (Fig. 2D). Additionally, the classical monocyte cluster cMo3 trended towards an increased abundance in both moderate and severe, compared to mild disease (Fig. 2D). Classical monocytes may contribute to the inflammation associated with COVID-19 severity, as they broadly upregulate activation markers during SARS-CoV-2 infection (*14*). In comparing markers defining the classical monocyte subsets (Fig. 2B and fig. S3), we observed that cMo1 and cMo2 have high CD33 expression. cMo1 also express CD45RA and CD163. cMo2 express CD9, HLA-DR, CD93, and CD36. cMo3 have moderate expression of CD33, CD36, and much lower expression of HLA-DR than do cMo1 and cMo2. Although we and others (*30*) observe an increase in classical monocytes amongst COVID-19 patients, the cMo1, cMo3 subsets specifically increased, whereas cMo2 decreased, with disease severity (Fig. 2D). cMo3 was the largest classical monocyte cluster in terms of cell frequency, and cMo4 was the smallest (Fig. 2A/D). cMo4 was abundant in several moderately and severely affected patients and was characterized by low expression of the siglec CD33, which is variably expressed in human PBMCs (*31*). CD56^+^ monocytes and CD3^+^ monocytes also exhibited slight reductions with disease severity (Fig. 2D).

**Figure 2.**
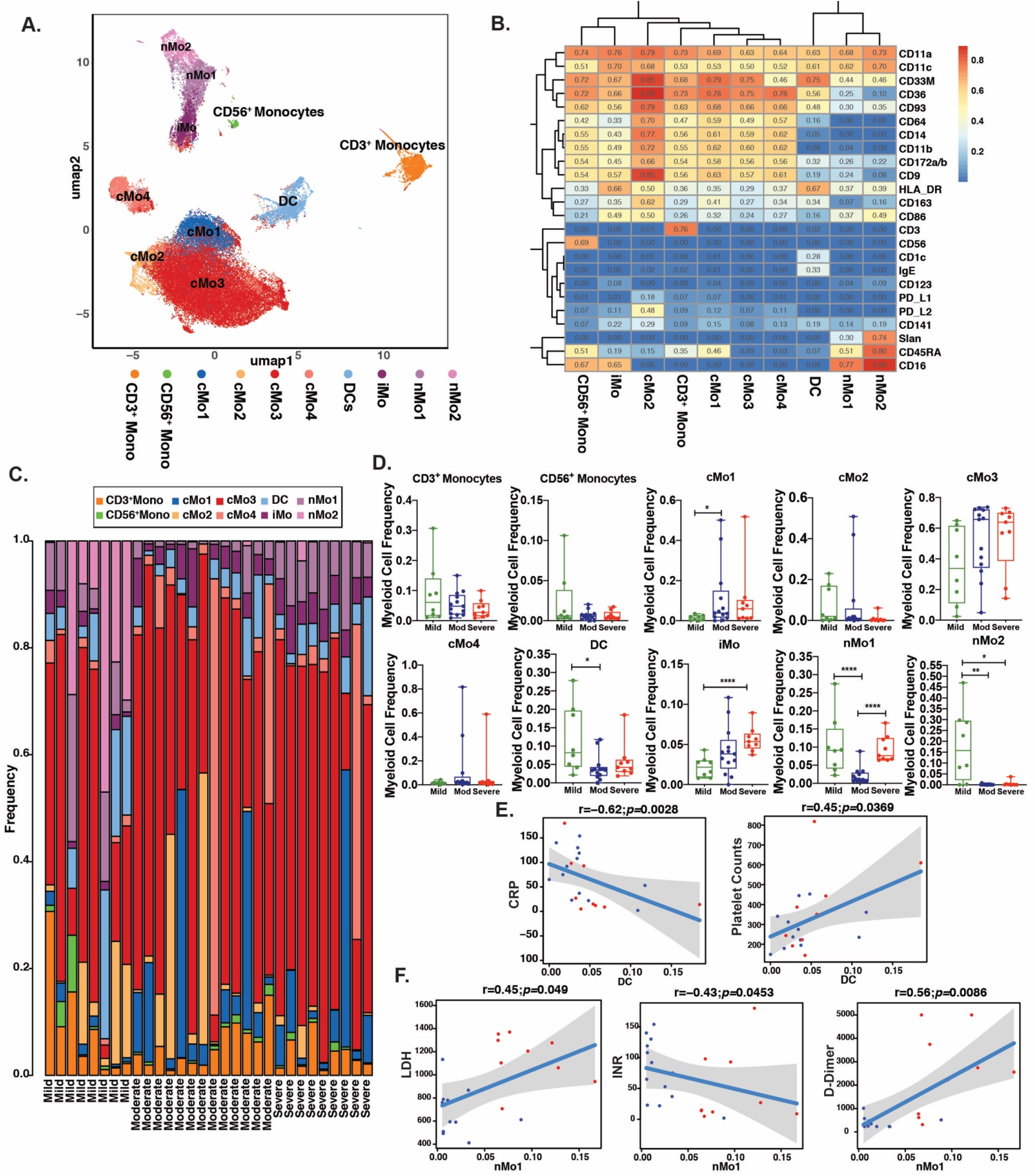
Monocyte immunoprofiing within recovered COVID-19 individuals. Myeloid cells were clustered and projected onto a UMAP, revealing 10 myeloid subsets (**A**). Heatmap of the median intensity of 24 markers across the 10 myeloid subsets (**B**). Stacked barplots of myeloid subset frequencies for each individual (**C**). Plotted frequencies of CD3^+^ monocytes, CD56^+^ monocytes, cMo1, cMo2, cMo3, cMo4, DC, iMo, nMo1, and nMo2 (**D**) for mild (n=9), moderate (n=13), and severe (n=9) individuals. Spearman correlation plots of DCs with CRP and Platelet counts for moderate (blue) and severe (red) individuals (E). Spearman correlation plots of nMo1 with LDH, INR, and D-Dimerfor moderate (blue) and severe (red) individuals (F). Statistical significance was determined using Wilcoxon rank sum test, “**p<0.001, **p<0.01, *p<0.05.

### Intermediate and Nonclassical Monocytes are Greatly Impacted by COVID-19

The most robust changes observed for myeloid cells in our cohort were those impacting the intermediate and nonclassical monocyte subsets. Intermediate monocytes (iMo) are CD14^+^CD16^+^ and elevated in atherosclerosis (*32*), linked to microinflammation in the vasculature in Type 2 diabetes (*33*), and associated with impaired vascular function in kidney disease (*34*). iMo are believed to represent a transitional monocyte stage through which classical CD14^+^ monocytes convert into nonclassical CD14^dim^CD16^+^ monocytes (*35*). Upon activation, classical monocytes are biased towards loss and gain of CD14^+^ and CD16^+^ expression, respectively (*36*). iMo cells express genes related to antigen presentation and participate in activation of the adaptive immune response, inflammation, and angiogenesis (*37*). In our COVID-19 cohort, the iMo cluster positively correlated with disease severity (p<0.05), with a 2.5-fold increase in iMo frequencies in mildly compared to severely ill individuals (Fig. 2D). An increase in pro-inflammatory iMo that expressed both IL-6 and GM-CSF was also reported in hospitalized COVID-19 patients in Hefei, China (*38*).

In contrast, CD14^dim^CD16^+^ nonclassical monocytes (nMo) play critical roles in maintaining vascular homeostasis (*39*) and guard against viral pathogens (*40*). Shulte-Schrepping et al. (*14*) recently reported depletion of all nonclassical monocytes in COVID-19. However, we know from our prior studies in human cardiovascular disease that nonclassical monocytes consist of distinct subsets (*27*). Here, in our COVID-19 cohort, nonclassical monocytes clustered into two separate populations (Fig. 2A). One population (nMo2) was positive for 6-sulfo-Lac-Nac (Slan), a glycosylated form of P-selectin Glycoprotein-1 (PSGL-1) (Fig. 2B). Slan^+^ nonclassical monocytes (called nMo2 here) have been identified in cancer (*41*), atherosclerosis (*27*), and multiple sclerosis (*42*), and were formerly designated as Slan-DCs (*43*). nMo2 produce IL-12 (*44*), IFNγ (*45*), activate both antigen-specific CD4^+^ and cytotoxic CD8^+^ T cells (*46, 47*), enable tumor phagocytosis and killing (*41, 48*), and enhance NK cell activation (*48, 49*). Thus, Slan^+^ nonclassical nMo2 monocytes regulate vascular homeostasis and drive immune responses. Here, we found that Slan^+^CD16^+^ nonclassical monocytes (nMo2) were nearly absent in both moderate and severe COVID-19 patients (Fig. 2C, light pink bars; Fig. 2D; *p* = 0.003 and *p*=0.01, Wilcox rank-sum test). Loss of nMo2 could thus promote progression to moderate and severe COVID-19 illness.

The other nonclassical monocyte subset, nMo1, was increased 4-fold in severely ill patients compared to moderately ill patients; still, this population was significantly lower in the moderate compared to the mild disease group (Fig. 2D). The cells within nMo1 lowly express Slan, but a previous study from our laboratory demonstrated that they do express CD40, CD63, CD93, and CD36 (*27*). The nMo1 cluster in moderately affected patients also exhibited higher expression of HLA-DR, CD45RA, PD-L1, and PD-L2 than that of mild or severe disease groups (Fig. 2B; fig. S3). CD45RA^+^ nonclassical monocytes have been associated with increased low density lipoprotein (LDL)-cholesterol levels (*29*). Interestingly, CD45RA is also a marker of myeloid progenitor cells, often found present in blood in acute myeloid leukemia patients (*50*), further supporting dysregulation of the homeostatic myeloid response by SARS-CoV-2 infection. The presence of PD-L1 and PD-L2 on the nMo1 cluster in moderately ill patients (compared to patients with mild or severe disease) suggests that these cells are unable to mount an effective immune response and may contribute to T cell exhaustion. Expression of PD-L1 and PD-L2 on nMo1 is lower in severe patients, yet the mechanism for this reduction of PD-L1 and PD-L2 in severely ill patients is unknown. Finally, as we included only the DC markers CD11c, CD123, CD1c, and CD141 in our CyTOF panel, we only observed 1 DC subset (Fig. 2A). This DC subset was decreased in moderate and severe patients compared to mild disease (Fig. 2C) as expected and observed in other studies (*15, 18*).

### Monocyte Phenotypes Associated with Clinical Features of COVID-19

We next asked if any monocyte or DC clusters were significantly associated with the clinical features of COVID-19 patients. We correlated myeloid cell frequencies with their matched inflammatory and coagulation parameters (shown in Fig. 1), as well as with COVID-19 severity, using Spearman rank correlations. DCs were negatively correlated with CRP levels (r=-0.62, *p*=0.003) and positively correlated with platelet counts (r=0.45, *p*=0.036) upon patient arrival to hospital (Fig. 2E). Of the classical monocyte clusters, cMo1 was associated with albumin levels, but this did not reach statistical significance. Thus, classical monocytes, although mostly increased in disease severity, were not stringently linked to any clinical parameter measured in our patients. In contrast, iMo were tightly and positively associated with disease severity (Fig. 2D), but were not significantly linked with any specific clinical parameters. There was a minor negative association with oxygen saturation supporting association with disease severity, but this was not significant (r=-0.4, *p*=0.06; data not shown).

The most striking findings arose from our analyses related to nonclassical monocyte phenotypes. nMo1 frequencies were higher in severely ill patients (Fig. 2D), positively correlated to the inflammatory marker LDH (r=0.45, *p*=0.04) (Fig. 2F), negatively correlated to levels of the coagulation indicator INR (r=0.54, *p*=0.009) (Fig. 2F), and positively correlated to D-dimers (r=0.56, *p*=0.008) (Fig. 2F) in these individuals. There was also a trend towards positive correlation of nMo1 with inflammatory CRP levels (r=0.37, *p*=0.08, data not shown). These data indicate that nMo1 levels are tightly associated with clinical parameters of coagulation and inflammation. Unfortunately, numbers of nMo2 were too low to power the assessment of associations of this cell type with disease severity.

### T Cell Heterogeneity in COVID-19 Patients

We next examined perturbations in the T cell compartment with disease severity within our cohort. CD8^+^ T cells and CD4^+^ T cells were re-clustered and projected onto a UMAP to obtain a detailed look at T cell heterogeneity (Fig. 3A). Using FlowSOM clustering and meta-clustering for merging subsets, we identified 17 clusters of T cells in PBMCs from our COVID-19 cohort. Specifically, we identified 7 CD4^+^ T cell subsets, 7 CD8^+^ T cell subsets, a subset of CD3^+^ T cells that were negative for CD4 and CD8 (double-negative (DN), CD4^+^CD8^+^ T cells (double-positive, DP), and CD3^+^CD56^+^ T cells (Fig. 3A). Within the CD4^+^ T cell compartment, 7 major cell clusters emerged on the basis of CD45RA, CD45RO, CCR7, CXCR3, CD11a, CD73, CD95, and PD-1 (Fig. 3B; fig. S4). In particular, we identified a CD4^+^ T naive (T_N_) (CD45RA^+^CCR7^+^CD45RO^lo^CD95^lo^) subset, a stem cell memory T (T_SCM_ - CD45RA^+^CCR7^+^CD45RO^lo^CD95^+^) subset, 2 central memory (T_CM_: CD45RA^lo^CCR7^+^CD45RO^+^) subsets, 2 effector memory (T_EM_: CD45RA^lo^CCR7^lo^CD45RO^+^) subsets, and a terminally differentiated effector (T_EMRA_: CD45RA^+^CCR7^-^CD45RO^lo^) subset. The 2 T_CM_ subsets were distinguished by low and high expression of the inflammatory chemokine receptor CXCR3. As the receptor for CXCL9, CXCL10, and CXCL11, CXCR3 is highly expressed within effector and memory subsets but largely absent on CD4^+^ T_N_ cells (*51*). CD4^+^ T helper 1 (Th1) cells have robust expression of CXCR3 (Fig. 3B) (*52*). Thus, we surmise that the CXCR3^+^ T_CM_ subset (T_CM_ 1) is Th1-like. The 2 remaining CD4^+^ T_EM_ subsets were distinguished by high and low PD-1 expression (Fig. 3B). PD-1 is associated with T cell exhaustion, but also upregulated on T cells that have recently encountered cognate antigen.

**Figure 3.**
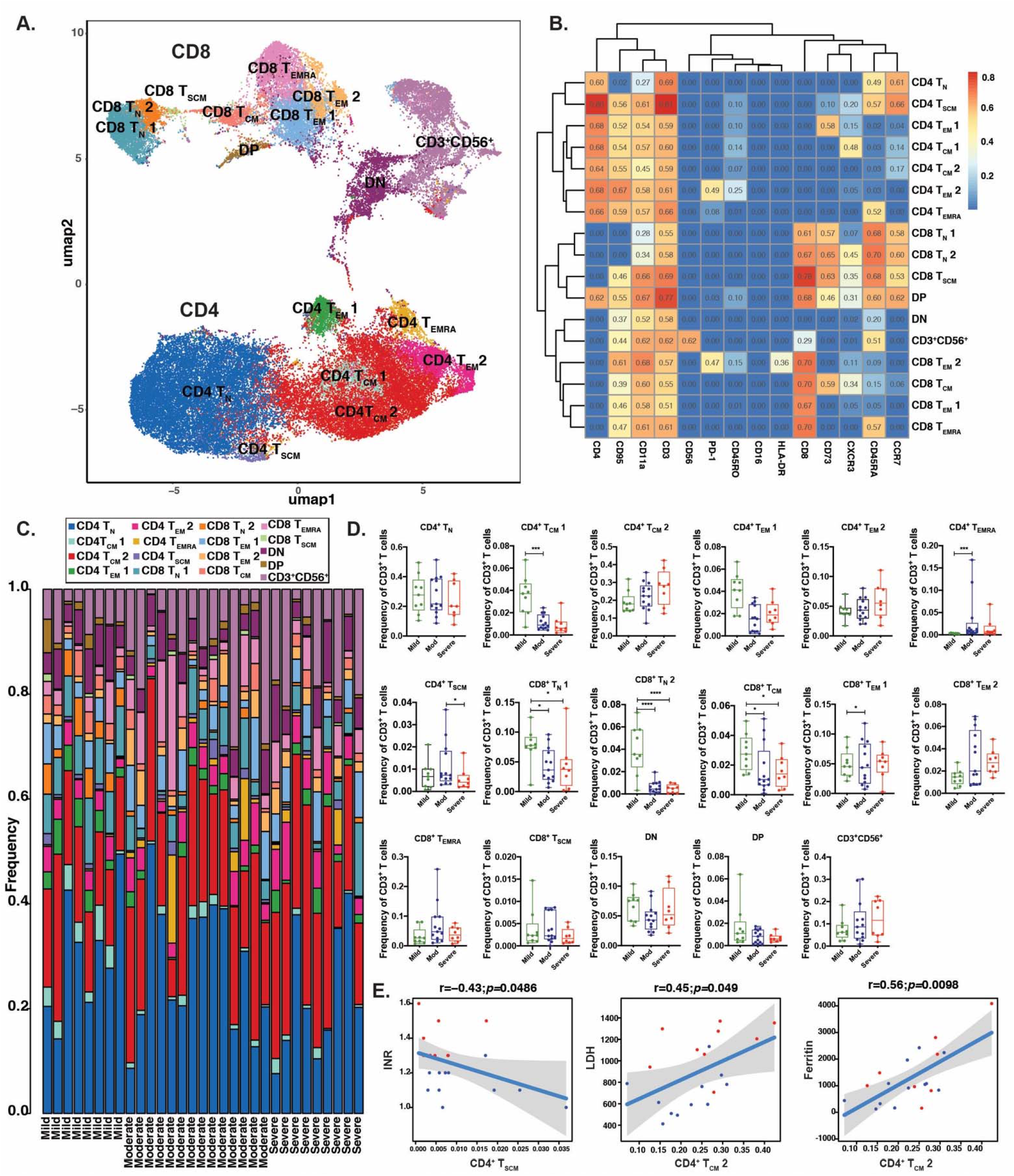
T cell immunoprofiiπg within recovered COVID-19 individuals. CD4^+^ and CD8^+^ **T** cells were clustered and projected onto a UMAP, revealing 17 T cell subsets (**A**). Heatmap ot the median intensity of 24 markers across the 17 subsets (**B**). Stacked barplots of T cell frequencies across mild, moderate, and severe individuals (**C**). Plotted frequencies of the 17 T cell subsets (**D**) for mild (π=8), moderate (n=13), and severe (n=9) individuals. Spearman correlation plots of CD4^+^ T_SCM_ cells with INR and CD4^+^ T_CM_ 2 cells with LDH and Ferritin (E). Statistical significance was determined using Wilcoxon rank sum test, ****p<0.001, **p<0.01, *p<0.05.

Within the CD8^+^ T cell compartment, we identified 2 CD8^+^ T_N_ subsets, a CD8^+^ T_SCM_ subset, 1 CD8^+^ T_CM_-like subset, 2 CD8^+^ TEM subsets, and 1 CD8^+^ T_EMRA_ subset (Fig. 3B; fig. S4). The 2 CD8^+^ TN subsets were distinguished by the pro-inflammatory chemokine receptor CXCR3. CD8^+^ T_N_ cells expressing CXCR3 possess increased effector potential compared to TN cells (*53*). Importantly, these cells are distinct from CD8^+^ T_SCM_ cells, as they lack CD95 expression. The CD8^+^ T_SCM_ subset expressed the T_N_ markers CD45RA, CCR7, and CD95. CXCR3 was also noted within this CD8^+^ T_SCM_ population and is a known marker of these cells (*54, 55*). The CD8^+^ T_CM_-like subset was CD45RA^lo^ and also expressed CCR7; however, expression of CD45RO was quite dim within the CD8^+^ T cell compartment overall (fig. S3). This CD8^+^ T_CM_-like cell was noted by high expression of the ecto-5’ nucleotidase CD73 and CXCR3. The CD8^+^ T_EM_ subset 2 was noted by high expression of the T cell exhaustion marker PD-1 and HLA-DR, whereas these markers were lowly expressed within CD8^+^ T_EM_ 1. Similar to the CD8^+^ T_CM_-like subset, expression of CD45RO was dim with CD8^+^ TEM 1, but CD45RA and CCR7 were not expressed. Analysis of subset frequencies revealed that these 17 subsets were expressed within all mildly, moderately, and severely ill individuals (Fig. 3C).

### Changes in T Cell Subsets with COVID-19 Severity

When quantifying CD4^+^ T cell subset frequencies in all patient groups, we observed significantly attenuated (3-fold) frequencies of the CD4^+^ T_CM_ 1 subset associated with high CXCR3 expression within mildly compared to moderately ill individuals (Fig. 3D). These T_CM_ cells, noted by high CXCR3 expression may embody a Th1-like subset. In anti-viral responses, Th1-polarized CD4 T cells propagate anti-viral CD8^+^ T cell responses. Our study indicates that this Th1 subset is lost in moderate disease. We also noted a significant (9-fold) enrichment in the CD4^+^ T_EMRA_ cell subset within moderately compared to mildly ill individuals (Fig. 3D). A recent study identified a significant increase in CD4^+^ T_EMRA_ frequencies in COVID-19 individuals compared to age-matched healthy controls; however, COVID-19 subjects in this study were not stratified by disease severity (*26*). Thus, moderately ill COVID-19 individuals display a loss of CD4^+^ CXCR3^hi^ T_CM_ cells and increased T_EMRA_ cells.

Examination of the CD8^+^ T cell compartment revealed 1.75-fold attenuations in the CXCR3^lo^ CD8^+^ T_N_ cell subset 1 in mildly compared to moderately ill and mildly compared to severely ill individuals (Fig. 3D). Moreover, the more effector-like CXCR3^+^CD8^+^ T_N_ subset 2 was decreased (6.6 and 8.8-fold, respectively) within mildly compared to moderately and severely affected individuals, respectively (Fig. 3D). The CD8^+^ T_CM_-like subset was significantly decreased by 1.7- and 1.8-fold within mildly compared to moderately and mildly compared to severely affected individuals, respectively (Fig. 3D). Moreover, the CD8^+^ T_EM_ 1 subset was increased within mildly compared to moderately ill individuals. Thus, severely ill COVID-19 individuals displayed enriched CD8^+^ T_EM_/T_EMRA_ populations and diminished TN subsets.

Analysis of our cohort (see Methods) confirmed that the mean age of non-hospitalized individuals with mild disease was significantly lower than that of the moderately-severely ill patients (\*\*\**p*=0.0004 (mild vs. moderate), \**p*=0.0161 (mild vs. severe), one-way ANOVA). As age is linked to T_N_ and T memory cell frequencies (*56, 57*), we decided to focus this study only on changes observed in the hospitalized patients (moderately + severely ill individuals), who were similar in age. We observed a 2-fold reduction in circulating CD4^+^ TSCM cells in moderate compared to severe COVID-19 disease (Fig. 3D). Human T_SCM_ cells, the least differentiated memory subset, are phenotyped as T_N_ cells that express CD95. T_SCM_ cells are highly proliferative, self-renewing, cytokine-producing, and endowed with the capacity to differentiate into T_CM_ and TEM subsets (*58, 59*). Importantly, within the T cell compartment, this CD4^+^ T_SCM_ cell cluster was the sole differentially-expressed subset between moderate and severe disease (Fig. 3D). Interestingly, while not statistically significant, CD8^+^ T_SCM_ cell frequencies were also similarly reduced in moderately versus severely ill COVID-19 individuals (Fig. 3D).

### T Cell Phenotypes Associated with Clinical Parameters of COVID-19

When correlated with clinical parameters, we observed a strong negative association of CD4^+^ TSCM frequencies from mildly and moderately affected individuals with international normalized ratio (INR) calculated from prothrombin time (r=-0.43, *p*=0.0488), which was significantly reduced in severe disease (Fig. 1J; Fig. 3E). Conversely, CD4^+^ T_CM_ 2 (CXCR3^lo^) frequencies were positively correlated with the inflammatory parameters lactate dehydrogenase (LDH; r=0.45, *p*=0.05) and ferritin (r=0.56, *p*=0.01) (Fig 3E). CD4^+^ T_EMRA_ frequencies were positively correlated with hemoglobin (r=0.45, *p*=0.04) (Fig. S5A). Examination of the CD8^+^ T cell compartment revealed that CD8^+^ T_SCM_ cell frequencies were inversely correlated with INR (r=-0.53, *p*=0.0016; Fig. S5B), while CD8^+^ T_EMRA_ frequencies were positively correlated with hemoglobin (r=0.46, *p*=0.03) (Fig. S5C) and albumin (r=0.44, *p*=0.0382; Fig. S5D).

### Interplay of Monocytes and T Cells in COVID-19 Severity

We next asked if the frequencies of monocyte subsets were associated with frequencies of specific T cells and if these correlations were linked to disease severity. While classical monocytes were not strongly linked with specific clinical measurements in our cohort, we did find that cMo3 frequencies (only) showed statistically significant associations with T cell subsets. In particular, we observed a negative association of cMo3 with CD4^+^ T_N_ cells (r=-0.51, *p*=0.0048; Fig. 4A). The cMo3 cluster was also significantly associated with the following T cell subsets that were changed with COVID-19 severity or that correlated with inflammatory indicators: CD8^+^ T_N_1 cells, CD4^+^ T_SCM_, as well as, CD4^+^ T_CM_2, CD4^+^ T_EM_2, and CD8^+^ T_EM_2. Specifically, the cMo3 cluster was negatively associated with CD8^+^ T_N_ 1 (CXCR3^lo^) and CD4^+^ T_SCM_ cells (Fig. 4A). In contrast to negative associations with T_N_ and T_SCM_, cMo3 was positively associated with CD4^+^ T_CM_ 2 (r=0.63, *p*=3e-4), CD4^+^ T_EM_ 2 (r=0.41. *p*=0.0268), and CD8^+^ T_EM_ 2 (r=0.46, *p*=0.011) (Fig. 4A). CD4^+^ T_EM_ 2 and CD8^+^ T_EM_ 2 may represent “exhausted” T cells, as these subsets were noted by PD-1 expression (Fig. 3B). A recent study found that stimulation of COVID-19 PBMCs elicited a profound CD4^+^ T_CM_ and a CD8^+^ T_EM_/T_EMRA_ response (*60*). Thus, cMo3 may contribute to immune dysregulation in response to COVID-19.

**Figure 4.**
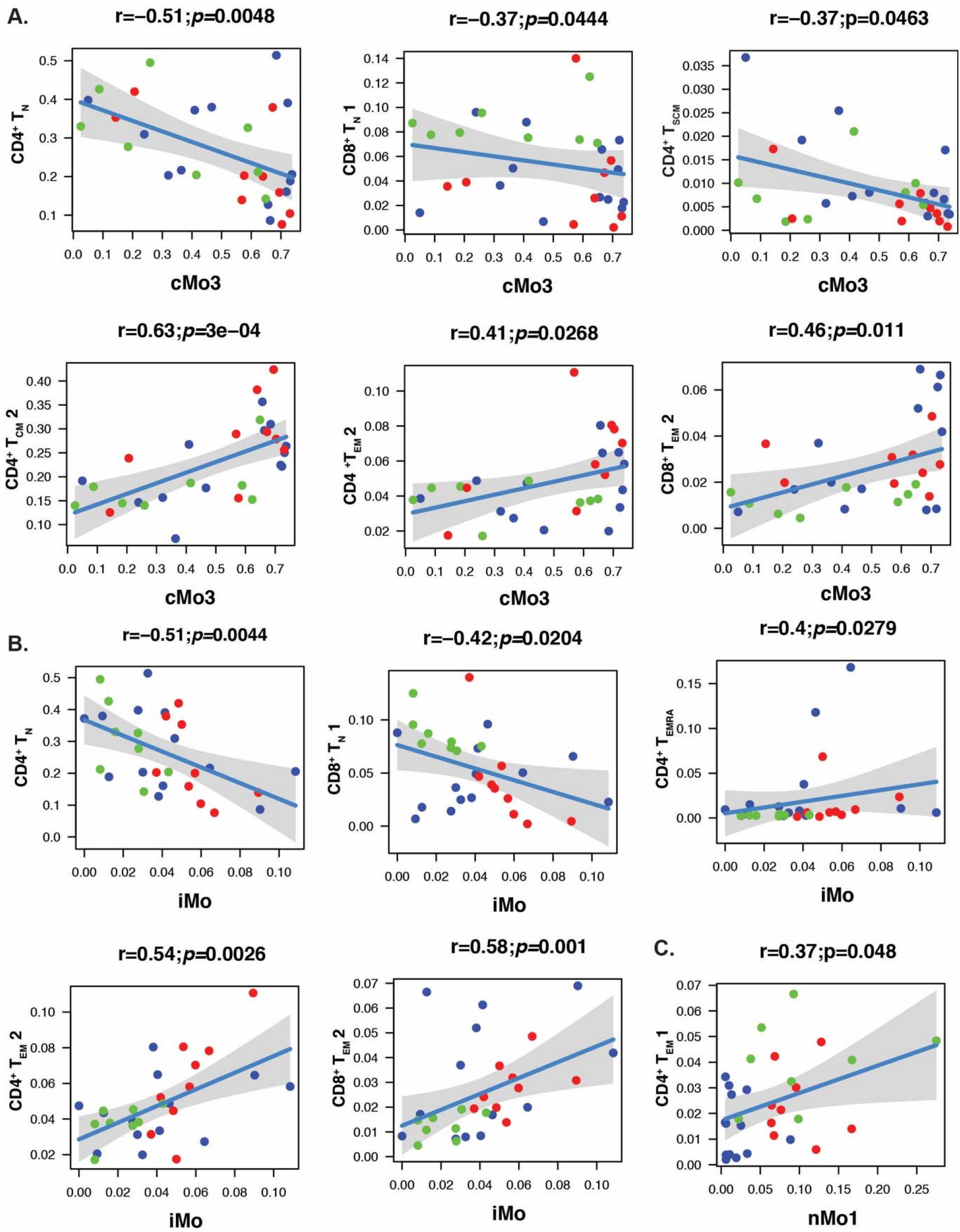
Monocyte and T cell subsets are correlated in COVID-19 individuals. Spearman correlation plots of cMo3 with CD4^+^ T_N_, CD8^+^ T_N_ 1, CD4^+^ T_CM_, 2, CD4^+^ T_EM_ 2, CD8^+^ T_EM_ 2 and CD4^+^ T_SCM_ (A), ÍMo with CD4^+^ T_N_, CD8^+^ T_N_ 1, CD4^+^ T_∈MRA_, CD4^+^T_EM_ 2, and CD8^+^T_EM_ 2 (B), and ⋂Mo1 with CD4^+^T_EM_ 1 (C) within mild (n=8, light green), moderate (n=l3, blue), and severe (n=9, red) individuals.

Additionally, we found that intermediate monocytes (iMo) showed the strongest correlation with T cell subsets that delineated disease severity (Fig. 4B). Specifically, iMo was negatively correlated with both CD4^+^ T_N_ (r=-0.51, *p*=0.0044) and CXCR3^lo^ CD8^+^ T_N_ (r=-0.42, *p*=0.0204) subsets, and positively correlated with corresponding effector CD4^+^ T_EMRA_ (r=0.4, *p*=0.0279), and the PD-1-expressing CD4^+^ T_EM_ 2 (r=0.54, *p*=0.0026), and CD8^+^ T_EM_ 2 subsets (r=0.58, *p*=0.001) (Fig. 4B). Cells in the iMo cluster express high levels of HLA-DR, and relatively high expression of CD9. CD9 is a tetraspanin present in plasma membrane microdomains, including lipid rafts (*61*). CD9 in monocytes and DC regulates HLA-DR trafficking, thereby influencing T cell activation (*61*). Thus, the high expression of HLA-DR in iMo may drive antigen presentation, and ultimately contribute to T cell exhaustion in these patients with severe illness. As cells within iMo are considered a transitional state in monocyte development, and because we found that iMo frequencies are significantly associated with COVID-19 disease severity (Fig. 2D), the correlation of iMo with these T cell subsets indicates potential likely functional crosstalk between fluid monocyte states and T cells during different stages of illness severity.

The nMo1 cluster was positively correlated with CD4^+^ T_EM_ 1 cells (r=0.37, *p*=0.0048) (Fig. 4C). These effector memory CD4^+^ T cells displayed high expression of CD73 (5’-ectonucleotidase), an ectoenzyme that catalyzes the conversion of 5’AMP to immunosuppressive adenosine (*62*). CD73 is a marker of T regulatory (Treg) cells; CD25^high^CD127^lo^ CD4^+^ T cells that express the forkhead box transcription factor Foxp3 and the ectoenzyme CD39 (*63*). Thus, this CD4^+^ TEM 1 subset is likely a Treg subset. However, as we didn’t include the markers CD25, CD127, Foxp3, or CD39 in our CyTOF panel, we cannot fully define this subset. CD4^+^ TEM 1 cells displayed trending reductions in frequency within mild compared to moderate and severe disease. In another study, Treg cells were attenuated in individuals with severe compared to mild COVID-19 (*64*). Similarly, our collaborators have found that antigenspecific SARS-COV2 reactive Treg cells are severely attenuated within hospitalized compared to nonhospitalized individuals (*65*). While lung tumor-infiltrating non-classical monocytes secrete chemokines that recruit T cells (*48*), the interplay between non-classical monocytes and Tregs is not well described. Further study is needed to determine the interplay between non-classical monocytes and Tregs during COVID-19.

Unfortunately, we were unable to perform correlations with either nMo2 or cMo4, due to their paucity in moderately and severely affected patients. In lieu of this method, we performed unsupervised clustering of PBMCs from all individuals by analyzing 487 combinations (10 myeloid subsets + 24 markers, 17 T cell subsets + 14 markers). Shown in Fig. S6 are the top differentially-expressed combinations of features across disease severity from this analysis. We found that CD141 expression in nMo2 (nMo2-CD14) and CD93 expression in both nMo1 and nMo2 (nMo2-CD93 and nMo1-CD93) best distinguished mildly ill patients (green bars) from the hospitalized patients (blue and red bars) (Fig. S6). Thus, the primary changes in the myeloid compartment that associate with COVID-19 severity lie in the nonclassical monocyte subsets. In addition, we observed a number of features that were dominantly expressed in moderately ill patients, such as PD1 expression in naive CD8^+^ (CD8^+^T_N_ 2-PD1) and CXCR3 in CD4^+^ T_CM_ (CD4^+^ T_CM_ 2-CXCR3) cells (Fig. S6). As a whole, we show that specific T cell and monocyte subsets are altered in the periphery of COVID-19 patients, highlighting systemic immune responses to the disease.

## DISCUSSION

In our current study, we examined myeloid and T cell heterogeneity in subjects recovering from mild, moderate, and severe COVID-19. Specifically, we identified 10 clusters of monocytes/DCs and 17 clusters of T lymphocytes via mass cytometry. Of these, clusters of intermediate and nonclassical monocytes, as well as CD4^+^ T_SCM_ cells, correlated with disease severity, coagulation factors, and inflammatory indices. We also found that intermediate monocytes tightly correlated with loss of T_N_ cells and an increased abundance of effector T cells, PD-1^+^ ‘exhausted’ CD4^+^ T_EM_ 2 cells, and CD8^+^ T_EM_ 2 cells. The nMo1 cluster also correlated with CD4^+^ T_EM_ 1 cells and disease severity.

The mildly ill individuals in our cohort were much younger than the moderately and severely ill individuals, while the latter two groups were similar in mean age (see Methods for details). This observation mirrored published findings that aged individuals are more susceptible to severe cases of COVID-19 (*66*), but is confounded by the fact that the mildly ill individuals represented a specific demographic (healthcare workers). Still, many pathological states are associated with age, including cardiovascular disease and cancer. A recent study by our laboratory demonstrated that human monocyte frequencies are minimally impacted by age (*27*). In contrast, naive T cell subsets markedly decrease with age and this change is accompanied by a concomitant increase in memory T cells (*56, 67, 68*). CD4^+^ TSCM cells embodied the sole T cell population found to be significantly attenuated in moderate versus severe disease. Moreover, TSCM cell frequencies were negatively correlated with the coagulation parameter INR in COVID-19 individuals. TSCM cells are antigen-experienced and the least-differentiated memory subset. In humans, T_SCM_ cells are distinguished from T_N_ cells by expression of CD95 (Fas or Apo-1). Increased TSCM cell frequencies are associated with improved response to viral infections. For example, HIV^+^ individuals on highly active antiretroviral therapy display increased CD8^+^ TSCM frequencies that negatively correlate with viral loads and CD4^+^ T cell counts (*69*). A recent study found no differences in peripheral CD4^+^ or CD8^+^ TSCM frequencies between COVID-19 donors and age-matched controls (*26*). However, as these data were not stratified by illness severity, longitudinal quantification of T cell subsets during COVID-19 is still needed.

Nonclassical monocytes play important anti-tumoral and atheroprotective roles (*48, 70*) and are distinguished by their ability to patrol the vasculature (*71, 72*). Vascular patrolling by nonclassical monocytes occurs during both steady-state and inflammation and allows surveillance against invading pathogens and maintenance of vascular endothelial homeostasis (*73*). We reported that the vascular endothelium is activated in mice that have nonclassical monocytes unable to patrol the vasculature due to disruptions in integrin signaling (*39*). We hypothesize that both nMo1 and nMo2 monocytes patrol the vasculature, but this has yet to be tested. We have found that nMo2 lowly express CXCR6, are migratory towards CXCL16, and highly phagocytic (*27*). We and others have found that Slan^+^ nMo2 monocytes make IL-12 (*44*), recruit NK cells (*48*), and activate T cells (*46, 47*).We speculate that loss of nMo2 cells in moderate and severe patients could be one explanation for the coagulation issues that arise in SARS-CoV-2 infection, in part due to the lack of the presence of these cells to maintain vascular endothelial homeostasis. However, studies have also shown that monocytes can release tissue factor, which is important for coagulation (*74*), so this could be another mechanism by which monocyte changes could impact coagulation in the vasculature of COVID-19 patients. Unfortunately, tissue factor was not included as a CyTOF marker in this study. Thus, an important next step of this study is to examine how alterations in coagulation are driven by COVID-19-based changes in monocyte subpopulations.

As we and others have long-hypothesized that all nonclassical monocytes are anti-inflammatory and anti-viral (*27, 40, 73, 75*), we were surprised to find that nMo1 monocytes are decreased in moderate but increased in severe COVID-19 patients. In fact, nMo1 cells appear to drive illness severity in our cohort. nMo1 cells in moderate patients displayed higher HLA-DR, PD-L1, and PD-L2 expression than those in severe patients, suggesting a possible exhaustion phenotype, or that they trigger T cell exhaustion. We also found positive correlations between these cells and levels of CRP, D-dimers, and INR. We were unable to perform functional studies on these cells, due to low amounts of sample availability. In lieu of this, we examined differences in marker expression between nMo1 and nMo2 cells. Differentially-enriched markers between these subpopulations included CD9 and CD36. CD9 on monocytes drives M2-like alternative activation pathways, including release of IL-10 (*76*). However, what drives the association of nMo1 with disease severity is unknown.

Recent work by the Amit laboratory (*77*) suggests that monocytes and macrophages can either be infected by, or phagocytize, SARS-CoV-2 particles. While monocytes minimally express the SARS-CoV-2 receptor ACE-2, they express CD147 (also known as BSG), a potential receptor for the virus (*78*). Given the viral scavenging of nMo1 and nMo2 cells, nMo2 monocytes may eventually be killed by viral infection or by excess viral particle uptake. An alternative hypothesis is that activation of nMo1 cells by viral entry or viral particle uptake triggers responses by nMo1 monocytes in COVID-19 patients. Future studies should focus on testing these two hypotheses.

Our study utilizes PBMCs from convalescent COVID-19 patients to characterize immune cell responses to the disease. Examining monocytes and T cells in acute COVID-19, as well as performing longitudinal studies on samples from patients during illness onset and recovery, could further define immune cell interactions during COVID-19 progression. In summary, our findings detail previously unknown phenotypes of monocytes and T cells during COVID-19 and suggest interactions between subsets of these two cell types during the disease.

## MATERIALS AND METHODS

### Patient Samples

Ethical approval for this study was received from the Berkshire Research Ethics 20/SC/0155 and the Ethics committee of the La Jolla Institute for Immunology. Written, informed consent was obtained for all 30 subjects. Moderate (n = 13) and severe (n = 9) individuals were hospitalized in a teaching hospital in the south of England either in a general ward (moderate disease), or with treatment escalated to transfer to the intensive care unit (ICU) for patients with severe disease. SARS-CoV2 was confirmed by reverse transcriptase polymerase reaction (RT-PCR) or by detection of anti-spike protein antibodies performed in the clinical virology laboratory of the hospital. The mild cohort comprised eight health care workers who were not hospitalized, but COVID-19 diagnosis was confirmed via RT-PCR or serological evidence of SARS-CoV2 antibodies. All individuals supplied up to 80mL of blood for research studies. Clinical and demographic data were collected from patient records for hospitalized patients, including ancestry, comorbidities, blood results, drug intervention, radiological involvement, thrombotic events, microbiology, and virology results.

The mean age of mild COVID-19 individuals was 39+2 (26-50) years, moderate: 62+14 (39-82) years; Severe: 56+10 (33-65) years (Mild vs. Moderate: *\*\*\*p*=0.0004; Mild vs. Severe: *\*p*=0.0161; Moderate vs. Severe: *p*=0.4076; One-Way ANOVA). Within the mild cohort, 50% were male, and within the moderate cohort, 77% of individuals were male. In the severe cohort, 78% were male. The mild cohort comprised 7 White/British individuals, and 1 White/Other individual. The moderate cohort comprised 2 Indian, 9 White/British, 1 Black/British, and 1 White/Other participant. The severe cohort comprised 5 White/British, 1 Black/British, 2 Indian, and 1 White/Other participant.

The average hospital stay of moderate individuals was 7.8+3.2 days, while severe individuals were in the hospital for an average of 20+11.8 days (*\*\*\*p*=0.004; Mann-whitney test). Five of 9 individuals (55.5%) within the severe cohort were intubated. Six individuals reported previous respiratory airway issues, including 3 within the moderate cohort reporting history of asthma or chronic obstructive pulmonary disease (COPD), 1 within the severe cohort with asthma, and two moderately affected individuals with COPD. Four of nine (44%) individuals within the severe cohort reported hypertension, while 1 of 13 (7.7%) individuals within the moderate cohort reported hypertension (*p*=0.1159; Fisher’s exact test). Two individuals (2/13) within the moderate cohort reported a previous DVT/arrhythmia and myocardial infarction, respectively. Two of thirteen individuals within the moderate cohort were diagnosed with cancer (*p*=0.4935; Fisher’s exact test). Three of nine (33.3%) individuals within the severe cohort had diabetes, in contrast to none in the moderate cohort (Moderate vs. Severe; *p*=0.0545; Fisher’s exact test). All moderate and severe individuals received antibiotics during hospitalization. Two of nine severe individuals were participants of the remdesivir and REMCAP trials, respectively. All hospitalized individuals recovered and were discharged from the hospital.

### Sample processing

PBMCs were thawed from the recovered COVID-19 individuals, barcoded with CD45, and stained with a monocyte-T cell panel, as previously described (*27*). Mild, moderate, and severe COVID-19 individuals were assayed over a period of five distinct CyTOF runs (Round 1: 3 moderate and 2 severe disease, Round 2: 5 moderate and 2 severe disease, Round 3: 2 mild, 5 moderate, and 1 severe disease, Round 4: 4 severe disease, and Round 5: 6 mild disease). To identify batch effects, a technical control of healthy PBMCs was CD45-barcoded and spiked into each individual sample. Tuning was performed every 6hrs. Data was normalized using the Matlab-based NormalizerR2013a_Win64. Live cells were gated via DNA-1 by DNA-2 and were Cisplatin^lo^.

### CyTOF analysis - Debarcoding and batch correction

Each CyTOF sample was debarcoded from the original fcs file (with spike-in healthy PBMC samples as a control) using a deconvolution algorithm (*79*) implemented in the CATALYST Bioconductor package. We reevaluated the markers used for barcoding by plotting the density of corresponding markers in each of fcs files after debarcoding. First, each fcs file was normalized using an arcsinh transformation (cofactor=5). Then, we took advantage of the same spike-in healthy control for the batch correction, using a quantile normalization method for the pooled distribution of each batch (a pair of sample and spike-in control), as implemented in the function *normalizeBatch* from the CYDAR Bioconductor package (*80*). The density plot of all protein markers was plotted before and after normalization; representative correction plots of CD3 (A), CD4 (B), CD8 (C), and CD11a (E), CD14 (F), and CD16 (G) are shown (Fig. S7).

### CyTOF analysis - High dimensional reduction and clustering

Mass cytometry analysis was performed similarly to the pipeline described in our previous work (*27*). We used the FlowSOM clustering method (*21*), a self-organizing map (SOM) machine learning method, with default parameters (SOM grid 10×10 corresponds to 100 subclusters) to identify major cell-types from CD45^+^ live cells as well as to determine the heterogeneity of myeloid and T cell compartments. Major cell-types were identified using well-known lineage markers. Consensus clustering was used to justify the optimal number of clusters from k=2 to k=30 to evaluate the robustness of the number of clusters based on relatively decrease in area under the CDF (Cumulative Distribution Function) curve (*81*). The clusters were merged based on their similarity of their representative protein markers. Heatmaps of median protein expression were produced using the pheatmap R package (v0.2). UMAP deduction was done using the umap R package (v0.2.3.1), a wrapper for Python package ‘umap-learn’. For T cell clustering in Figure 3, CD14^lo^CD66b/Siglec/CD19^lo^ CD3^+^CD4^+^ and CD3^+^CD8^+^ T cells were gated in FlowJo v10.3 prior to downstream FlowSOM clustering. Differential expression analysis of combined features of Tcell/myeloid subsets and surface markers (487 total features) were used to select top features differentiating disease severity (fold-change cutoff of 1), as shown in Figure S6.

### Statistical analysis

Differences in immune subpopulation frequencies were evaluated using Wilcox rank-sum tests. Correlation of T cell and monocyte/DC subset frequencies with one other, as well as with clinical parameters, was performed using Spearman rank correlation tests.

## ACKNOWLEDGEMENTS

We would like to thank Cheryl Kim and the LJI Flow Cytometry Core for assistance. We also thank Luke Smith for patient recruitment and sample collection, Callum Dixon, Benjamin Johnson, Lydia Scarlett and Silvia Austin for collection of clinical data, and Céline Galloway, Oliver Wood, Katy McCann and Lindsey Chudley for sample processing.

## Author Contributions

Dr. Padgett designed and performed the experiments, generated figures, and wrote the manuscript. Dr. Dinh analyzed all CyTOF data and performed correlational analyses. Dr. Chee recruited and cared for patients and obtained blood samples. Dr. Olingy assisted with experimental design. Ms. Wu assisted with antibody conjugations and CyTOF analysis. Dr. Araujo participated in study design, contributed to writing and editing the manuscript, and participated in helpful discussions. Dr. Vijayanand participated in helpful discussions. Dr. Ottensmeier led the clinical study in the UK, edited the manuscript, and participated in helpful discussions. Dr. Hedrick directed the study and wrote the manuscript.

## Data and Materials Availability

Complete CyTOF panels used including metals and antibody clones are listed. All CyTOF data will be available at http://flowrepository.org/.

