## Supplementary Figures 1-7 for "Interplay of Monocytes and T Lymphocytes in COVID-19 Severity"

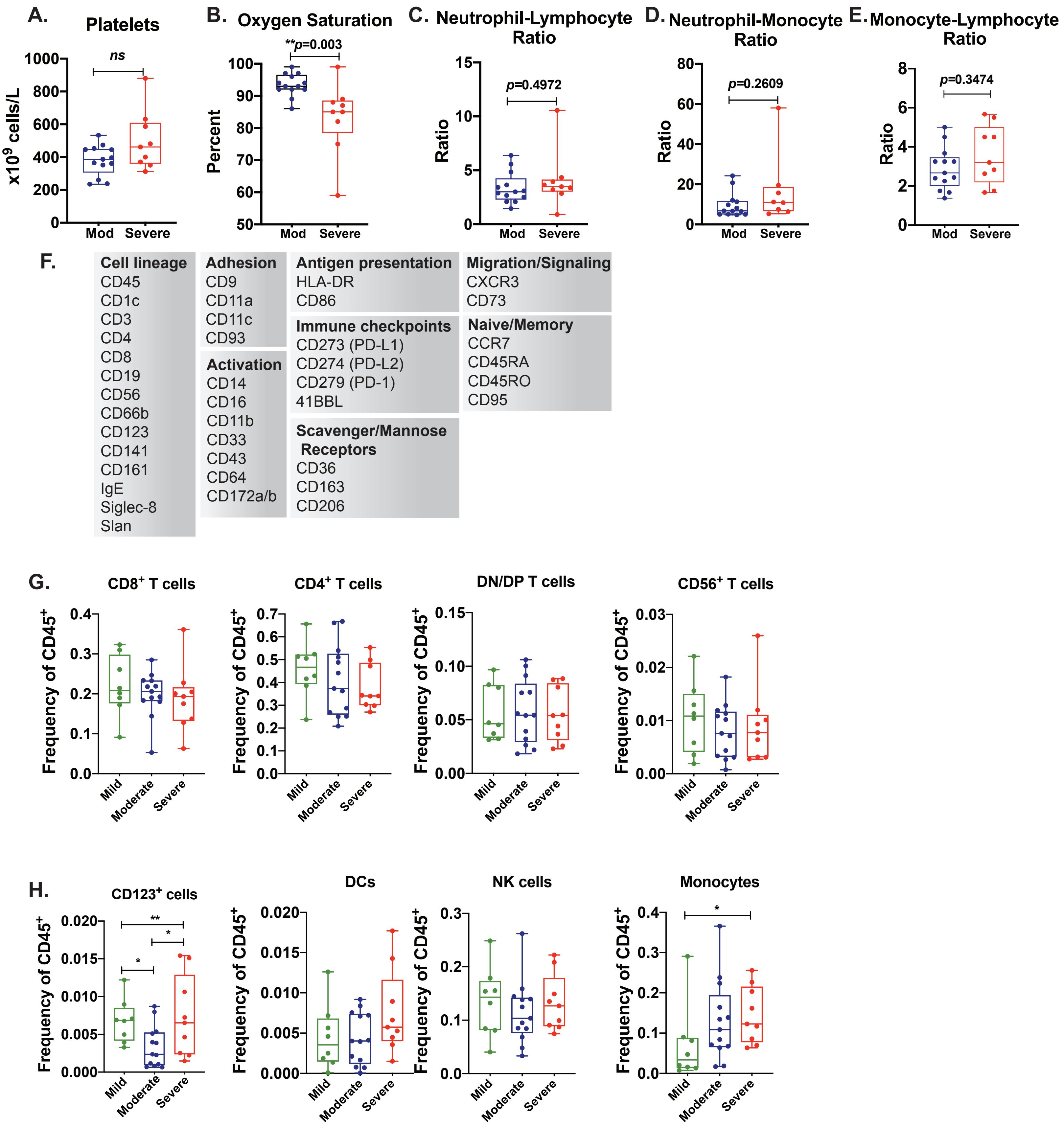


**Supplemental Figure 1. Patient parameters, CyTOF panel, and major immune cell population changes in recovered COVID-19 individuals.** Platelets (**A**) oxygen sauturation (**B**), neutrophil-lymphocyte ratio (**C**), neutrophil-monocyte ratio (**D**), monocyte-lymphocyte ratio (**E**). Panel for examining myeloid and T cell heterogeneity via CyTOF (**F**). Plotted frequencies of CD8$^+$ T cell, CD4$^+$ T cells, DN/DP T cells, CD56$^+$ T cells (**G**), CD123$^+$ cells, DCs, NK cells, and monocytes (**H**) in mild (n=8), moderate (n=13), and severe (n=9) individuals. Statistical significance was calculated using Mann-whitney test (**A-E**) and wilcoxon sum-rank test (**G-H**). *$p$<0.05

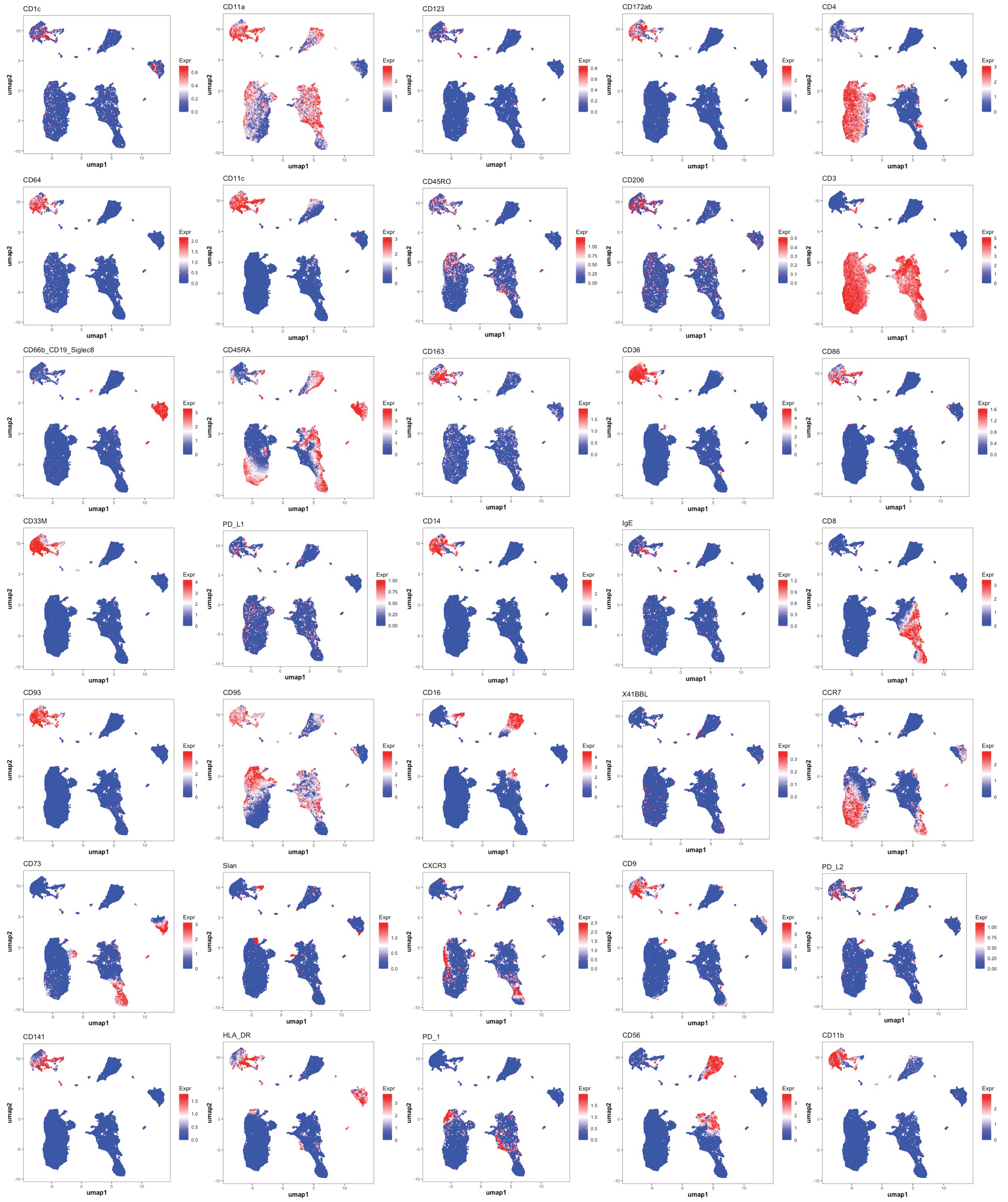


**Supplemental Figure 2. UMAP of CD45$^+$ leukocytes represented via color by channel.** CD45$^+$ leukocytes from COVID-19 individuals were clustered and projected onto a UMAP, with intensity of the markers utiized in the CyTOF panel represented via color by channel.

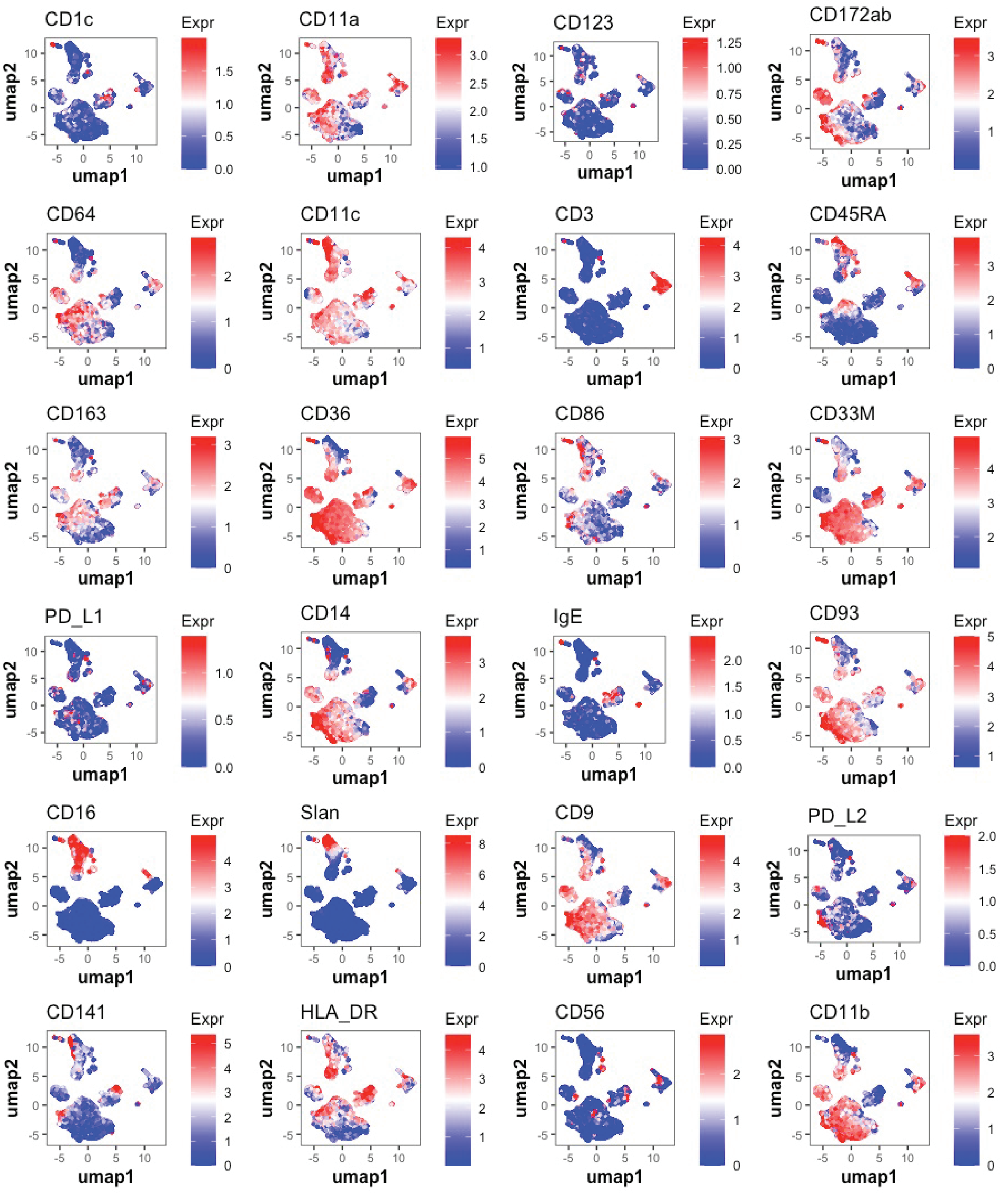


**Supplemental Figure 3. UMAP of myeloid compartment represented via color by channel.** Myeloid cells were clustered and projected onto a UMAP, with intensity of CD1c, CD11a, CD123, CD172a/b, CD64. CD11c, CD3, CD45RA, CD163, CD36, CD86, CD33M, PD-L1, CD14, IgE, CD93, CD16, Slan, CD9, PD-L2, CD141, HLA-DR, CD56, and CD11b represented via color by channel.

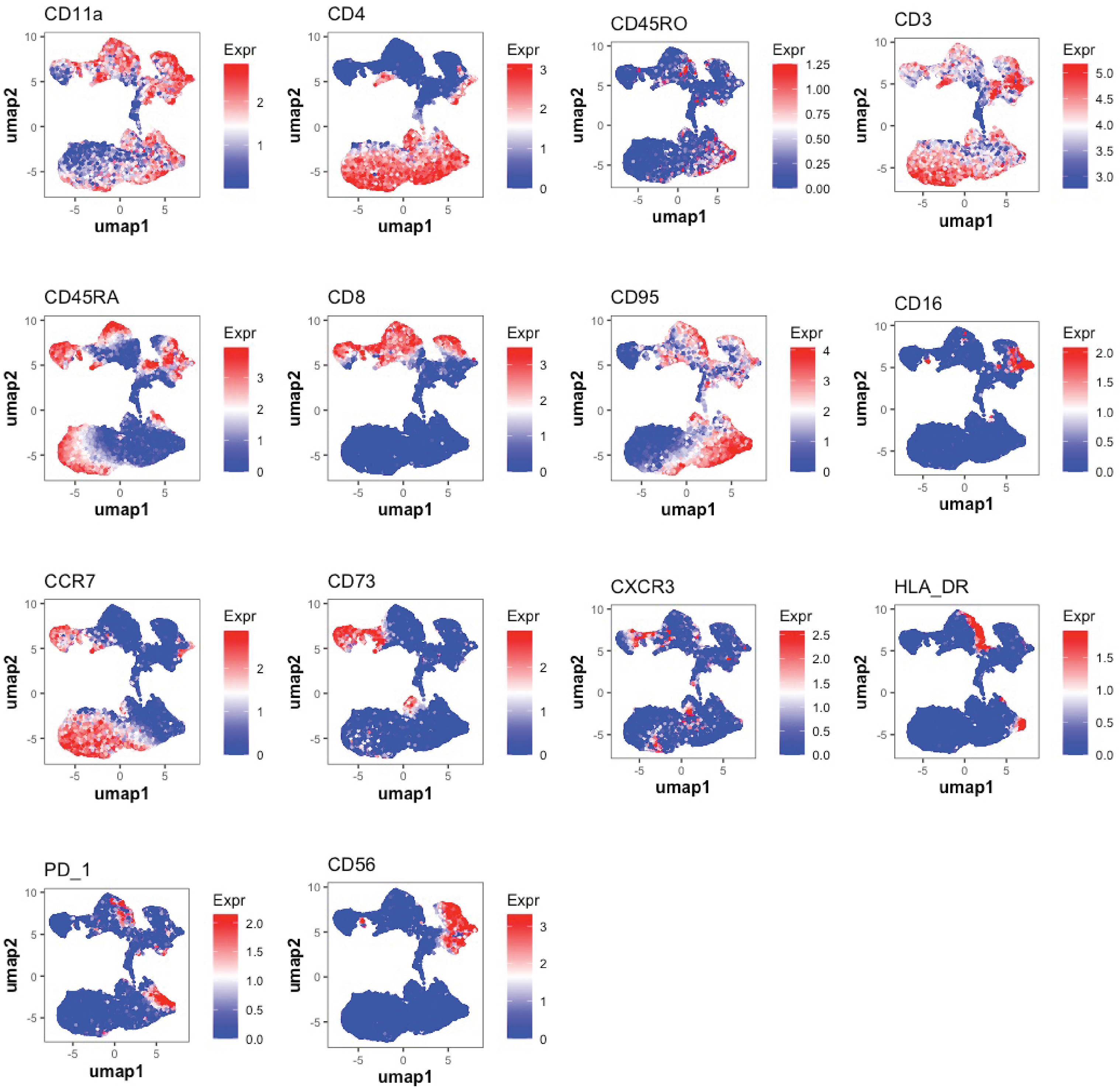


**Supplemental Figure 4. UMAP of T cell compartment represented via color by channel.** T cells were clustered and projected onto a UMAP, with intensity of CD11a, CD4, CD45RO, CD3, CD45RA, CD8, CD95, CD16, CCR7, CD73, CXCR3, HLA-DR, PD-1, and CD56 represented via color by channel.

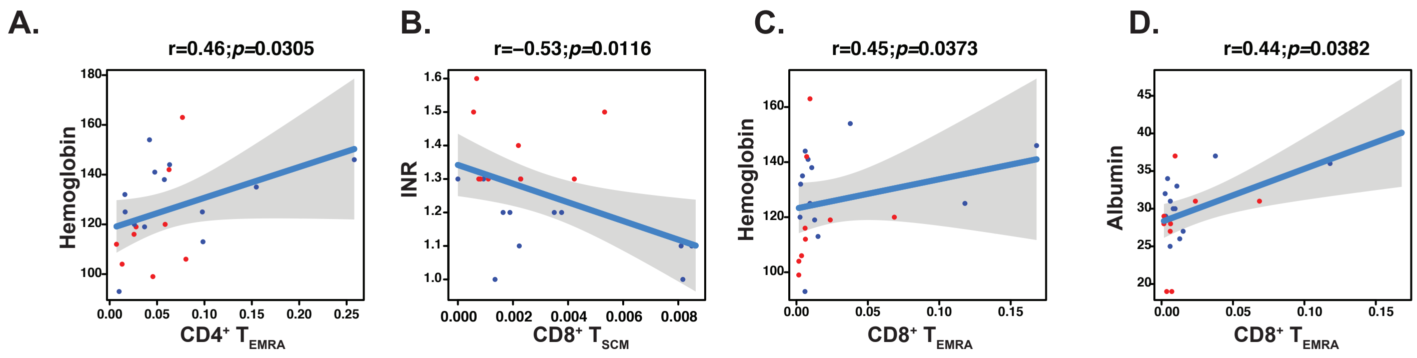


**Supplemental Figure 5. Spearman Correlation Plots for T cell subsets with COVID-19 clinical parameters.** Spearman correlation plots of CD4$^+$ T$_{EMRA}$ with Hemoglobin (**A**), CD8$^+$ T$_{SCM}$ cells with INR (**B**), CD8$^+$ T$_{EMRA}$ cells with hemoglobin (**C**) and albumin (**D**).

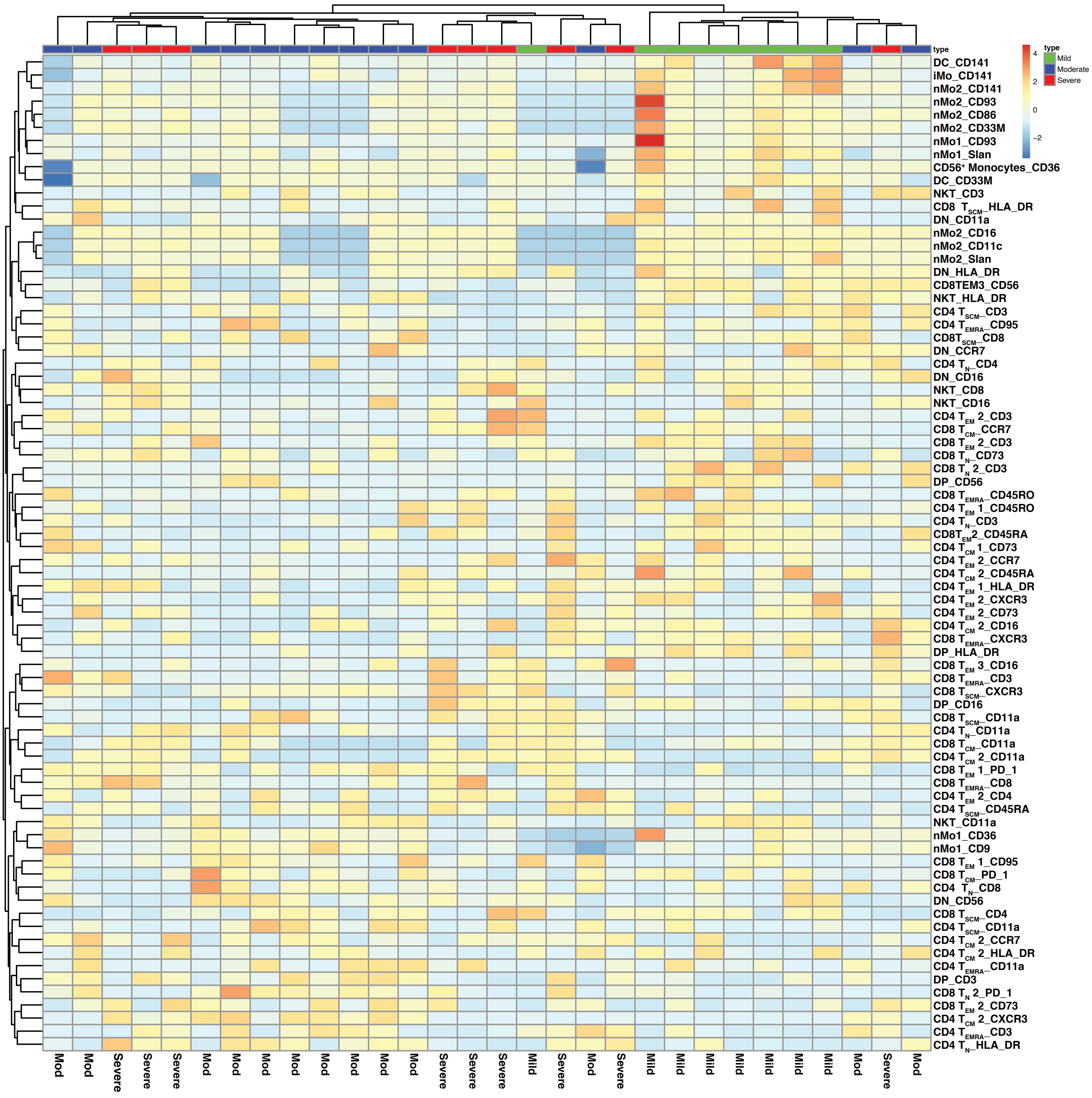


**Supplemental Figure 6. Unsupervised clustering of 30 recovered COVID individuals.** Unsupervised clustering of 30 recovered COVID samples based on the top differentially expressed combinations of marker and subset protein surface markers (76 out of 478 combinations) across disease severity. Samples annotated by disease severity are located on the x-axis, while the combinations of subset name and marker name are located on the y-axis. The median expression was normlized using z-score.

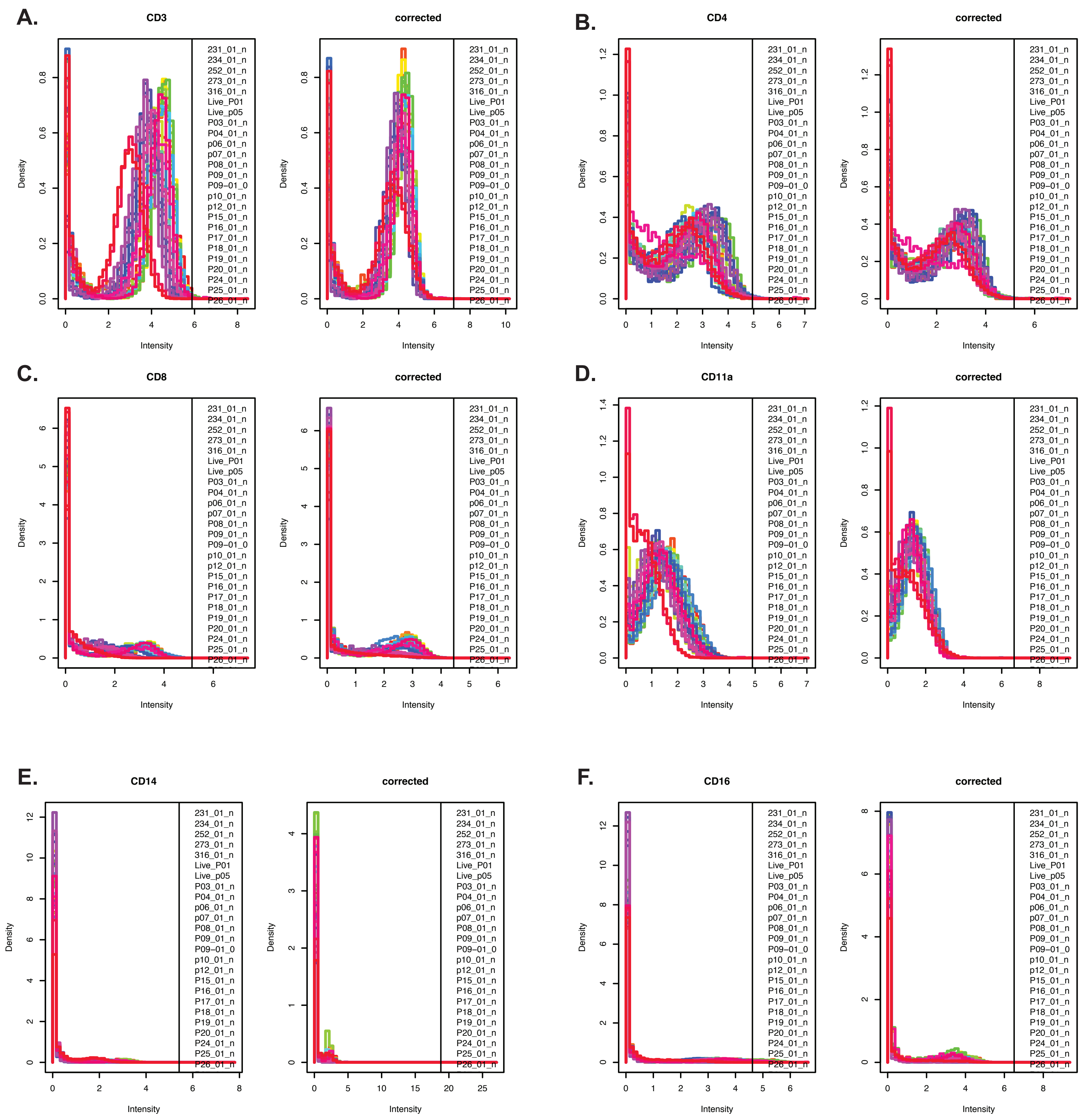


**Supplemental Figure 7. CyTOF batch correction analysis.** The density of protein markers CD3 (**A**), CD4 (**B**), CD8 (**C**), CD11a (**D**), CD14 (**E**), and CD16 (**F**) was plotted before and after batch correction.
